# Micro-CT analyses of the lung in mice: Parameters influencing the radiation dose and acquisition quality

**DOI:** 10.1101/2022.04.27.489643

**Authors:** Sandrina Körner, Christina Körbel, Yvonne Dzierma, Katharina Speicher, Matthias W. Laschke, Christian Rübe, Michael D. Menger, Maximilian Linxweiler

## Abstract

Microcomputed tomography (micro-CT) is a frequently used imaging tool for a wide spectrum of *in-vivo* mouse models in basic and translational research. To allow an accurate interpretation of micro-CT images, high spatial resolution is necessary. However, this may also lead to a high radiation exposure of the animals. Therefore, animal welfare requires exact information about the expected radiation doses for experimental planning. To gain this, a mouse cadaver was herein used for micro-CT analyses under different conditions. For each radiation dose measurement, the cadaver was labeled with thermoluminescent dosimeter chips around the thoracic skin surface. Micro-CT scans of the thorax were performed with spatial resolutions of 35 µm, 18 µm and 9 µm in combination with Al0.5, Al1.0, CuAl and Cu filters. As a surrogate of image quality, the number of identifiable lung vessels was counted on a transversal micro-CT slice. Measured radiation doses varied from 0.09 Gy up to 5.18 Gy dependent on resolution and filter settings. A significant dose reduction of > 75% was achieved by a Cu filter when compared to an Al0.5 filter. However, this resulted in a markedly reduced image quality and interpretability of microstructures due to higher radiation shielding and lower spatial resolution. Thus, the right combination of distinct filters and several scan protocol settings adjusted to the individual requirements can significantly reduce the radiation dose of micro-CT leading to a higher animal welfare standard.

## Introduction

In preclinical research, microcomputed tomography (micro-CT) has become one of the most commonly applied imaging modalities for studying *in-vivo* disease models ^1^. Rodents, particularly mice and rats, are used in most of these models. The micro-CT technology facilitates a longitudinal and non-invasive study design to implement the 3R principles in terms of reducing the number of animals and biological variability, because the analyzed animals can function as their own controls ^2,3^.

The non-invasive and quantitative investigation of disease onset, progression and/or therapeutic approaches in tumor studies still remains a major challenge in small animal CT-imaging ^4–6^. The assessment of tumor burden as well as the detection of metastases in mice require much higher spatial resolution, sensitivity and tissue contrast when compared to human CT imaging ^3^. To achieve this, specific refinements to the instrumentation have been introduced ^7^. Accordingly, micro-CT scanners currently used in preclinical imaging provide excellent three-dimensional (3D) images with a high spatial resolution (5-150 µm) on nearly a microscopic level ^8,9^. However, high image quality and resolution are at the expense of a markedly higher radiation exposure ^10–12^. While a conventional CT scan of the lung in humans exposes the patient to a radiation dose of ∼5 mSv, the dose of micro-CT scans in mice is at a ∼100-fold higher level. Hence, special attention should be given to the aspects of radiation protection in respect of the 3R principle, particularly in longitudinal studies, in which repeated micro-CT acquisitions are performed. In fact, even the exposure to a low radiation dose may have a strong impact on the immune response and other biological mechanisms determining the study outcome ^13^.

The manufacturers of commercially available preclinical micro-CT scanners provide only limited data for radiation exposure. Moreover, the radiation sensitivity markedly differs between different mouse strains ^14–16^. Thus, radiation dosimetry may represent a valuable tool in the planning of experimental animal studies but has rarely been investigated and not been standardized so far. Therefore, we have established an experimental setup for radiation dosimetry during micro-CT scans of the lung in mice. Our study illustrates the essential influence of different filters and spatial resolution on radiation exposure and image quality and provides a worthful methodological tool for a reasonable planning of imaging modalities in longitudinal animal studies.

## Materials and Methods

### TLD dosimetry and calibration

For dosimetry measurements during micro-CT image acquisition, an adult male mouse cadaver of the NOD-Scid strain (age 16 weeks) without pretreatment was used (Janvier Labs, Le Genest-Saint-Isle, France). The experiments were approved by the local governmental animal protection committee (permission number: 68/2015) in accordance with the German legislation on protection of animals, the EU Directive 2010/63/EU and the National Institutes of Health Guidelines for the Care and Use of Laboratory Animals (NIH Publication #85-23 Rev. 1985).

Dose measurements were performed using Harshaw thermoluminescent dosimeter chips TLD 100H which were read, heated and annealed in a Harshaw TLD 5500 reader (Thermo Fisher Scientific, Waltham, MA, USA) with the vendor-recommended time-temperature protocol (preheating at 145 °C for 5 seconds, acquisition with 10 °C/s temperature gradient to 260 °C held for 23 1/3 seconds and annealing at 260 °C for 20 seconds). Calibration of the TLDs was carried out at 1 Gy in a 6 MV photon beam from a Siemens Artiste (Siemens Healthcare, Erlangen, Germany) medical linear accelerator since an unrealistically long irradiation time would have been required for calibration in a ^90^Sr/^90^Y irradiator. For calibration, the TLDs were placed in an insert beneath 35 mm of acrylic glass slabs. In a second measurement, a PTW semiflex 31013 ionization chamber (PTW-Freiburg, Freiburg, Germany) was positioned at the same location so that the absolute dose at this point could be accurately verified, including air density, beam quality, and radius correction. Due to small deviations in dose within the photon beam (flatness and symmetry better than 3 %), deviations of the same order of magnitude are inherent in this calibration. To check the precision and reproducibility of the calibration, TLDs were subsequently irradiated several times with a dose of 1 Gy and compared with ionization chamber measurements. The average reading of the chip set always remained within ±5 % of the irradiated dose as measured by the ionization chamber.

As an independent check of the TLD calibration, the TLDs were exposed to 100 mGy in the ^90^Sr/^90^Y irradiator, which confirmed a measurement accuracy below 5 % ^17^. For lower photon energies, energy correction factors known from previous experiments were applied. TLD measurements at source-surface distance of 100 cm with 70 kV and 63 mAs compared with absolute dose as measured by a PTW DALI detector and 77334 ionization chamber (PTW-Freiburg, Germany), yielding a correction factor of 0.86 consistent with literature data ^18–20^. For our applications at 50 kV voltage, the relative response of the TLDs remained in the same range (6 % change in relative dose response between N-30 and N-100 ^19^) so that the same factor was applied to all readings.

### Dose measurements

The mouse cadaver was used as a phantom body for dose measurements. For each measurement protocol, TLDs were placed at four positions: ventral, left, dorsal and right position at the thoracic skin surface of the mouse (Figure 1). Three TLDs were placed at each position.

**Figure 1:**
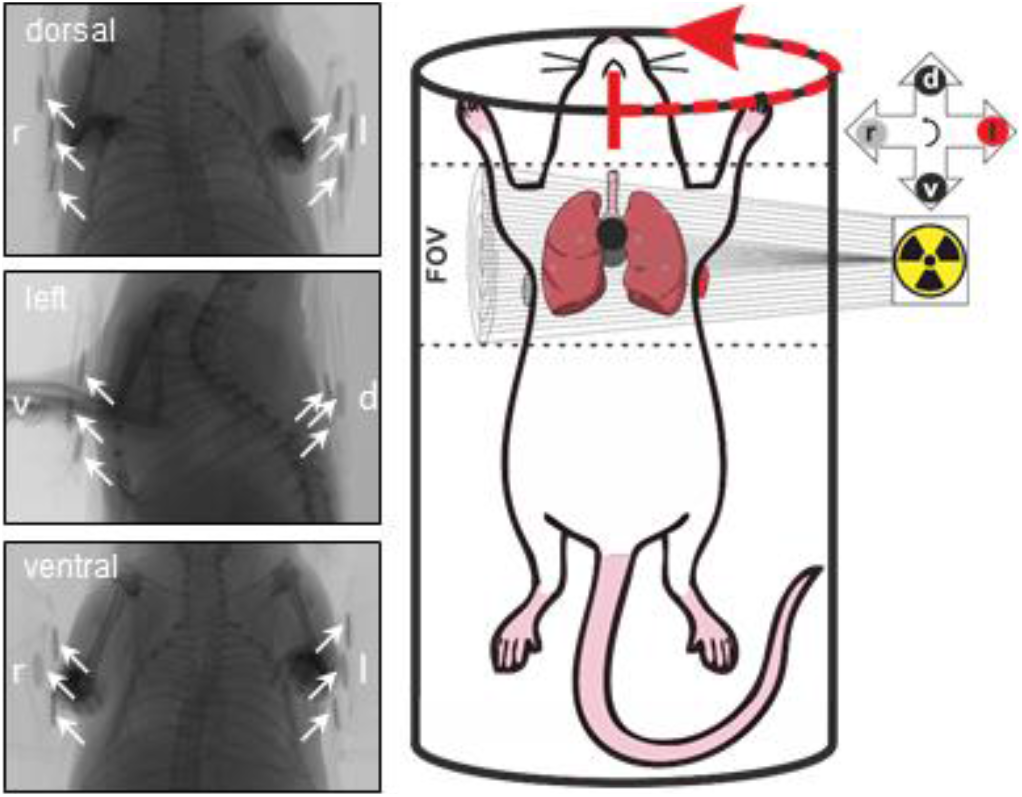
Experimental setup. For micro-CT imaging, a mouse cadaver was placed in supine position onto an animal bed. For each CT protocol and dose measurement, three TLDs each were placed in the following four positions: ventral (black), left (red), dorsal (black) and right (grey) position on the thorax of the cadaver. The field of view (FOV), in which the image acquisition was performed over 180° by 0.7° and 1.0° rotation steps, is marked with dashed lines. Counterclockwise rotation direction of the x-ray source is indicated. On the left x-ray images, the TLD locations are labeled with arrows.

After setting the field of view to the thoracic region of the mouse, micro-CT imaging was performed by recording X-ray pictures counterclockwise over 180 degrees (from ventral over left to dorsal position) with either 0.7° or 1.0° rotation steps (Figure 1) using a SkyScan1176 micro-CT scanner (Bruker AXS, Karlsruhe, Germany). For better image quality every X-ray picture was calculated out of two single recorded pictures. The filters in the X-rays changed from aluminum (Al) 0.5 mm (Al 0.5), Al 1 mm (Al 1), copper / aluminum (CuAl) to copper (Cu) filter in each experimental setup. The voltage of the micro-CT acquisition was 50 kV. The resolution of the micro-CT image depended on the current and exposure time so that a distinction could be made between a resolution of 35.28 µm (35 µm), 17.64 µm (18 µm), and 8.82 µm (9 µm). All micro-CT protocols differed from each other by the used filter, rotation step and resolution (pixel size). The voltage and frame averaging were identical for all measurements, the value for charge and the scan duration was dependent on the resolution.

In table 1 all different used micro-CT measurement settings were summarized and grouped together (group A, B, C, D).

**Table 1:** Micro-CT scanner settings for TLD measurement and image acquisition. Each measurement setup of different scan parameters is grouped together (A, B, C, D). Setting differences between single groups are highlighted.

| group | filter | rotation<br>step<br>[deg] | pixel<br>size<br>[μm] | Charge<br>[mAs] | (current x<br>exposure)<br>[μA] x [ms] | Voltage<br>[kV] | frame<br>averaging | scan<br>duration<br>[s] |
| --- | --- | --- | --- | --- | --- | --- | --- | --- |
| A1) | Al 0.5 | **180 / 0.7** | 35.28 | 0.03 | 476 x 58 | 50 | 2 | 254 |
| A2) | Al 1 | **180 / 0.7** | 35.28 | 0.03 | 476 x 58 | 50 | 2 | 255 |
| A3) | CuAl | **180 / 0.7** | 35.28 | 0.03 | 476 x 58 | 50 | 2 | 255 |
| A4) | Cu | **180 / 0.7** | 35.28 | 0.03 | 476 x 58 | 50 | 2 | 256 |
| B1) | Al 0.5 | **180 / 1.0** | **35.28** | 0.03 | 476 x 58 | 50 | 2 | 179 |
| B2) | Al 1 | **180 / 1.0** | **35.28** | 0.03 | 476 x 58 | 50 | 2 | 178 |
| B3) | CuAl | **180 / 1.0** | **35.28** | 0.03 | 476 x 58 | 50 | 2 | 179 |
| B4) | Cu | **180 / 1.0** | **35.28** | 0.03 | 476 x 58 | 50 | 2 | 180 |
| C1) | Al 0.5 | 180 / 1.0 | **17.64** | 0.11 | 500 x 220 | 50 | 2 | 351 |
| C2) | Al 1 | 180 / 1.0 | **17.64** | 0.11 | 500 x 220 | 50 | 2 | 351 |
| C3) | CuAl | 180 / 1.0 | **17.64** | 0.11 | 500 x 220 | 50 | 2 | 400 |
| C4) | Cu | 180 / 1.0 | **17.64** | 0.11 | 500 x 220 | 50 | 2 | 375 |
| D1) | Al 0.5 | 180 / 1.0 | **8.82** | 0.40 | 500 x 800 | 50 | 2 | 1190 |
| D2) | Al 1 | 180 / 1.0 | **8.82** | 0.40 | 500 x 800 | 50 | 2 | 1193 |
| D3) | CuAl | 180 / 1.0 | **8.82** | 0.40 | 500 x 800 | 50 | 2 | 1172 |
| D4) | Cu | 180 / 1.0 | **8.82** | 0.40 | 500 x 800 | 50 | 2 | 1131 |

After dosimetry, all micro-CT protocols were repeated in exactly the same position, regardless of the acquisition settings, which provided more comparable images for the image quality analysis (no dose measurements were performed at this point). Thereafter, the gray scale distribution was measured in the micro-CT images of the different protocols using the ImageJ 1.48v software (National Institutes of Health, USA).

### Statistics

Data were first analyzed for normal distribution and equal variance. Differences between the experimental groups were calculated by applying an ANOVA test followed by the appropriate post-hoc comparison. The post-hoc analyses included the correction of the α-error according to Bonferroni to compensate for multiple comparisons. In addition, a linear regression analysis was performed (Pearson’s coefficient of correlation for parametric distribution) to evaluate the correspondence of different micro-CT protocols (SigmaStat; Jandel Corporation, San Rafael, CA, USA). Data are given as mean ± standard error of the mean (SEM) or boxplots (median, 1^st^ and 3^rd^ quartile). Statistical significance was considered for p-values <0.05.

## Results

### Impact of the rotation steps on the radiation burden

In a first set of experiments, the radiation dose was measured around a mouse thorax while recording micro-CT images with Al 0.5, Al 1, CuAl, and Cu filters. During this process, images were recorded counterclockwise over 180° starting from ventral and running over the left thoracic side to the dorsal side. Two analyzed measurement settings differed in the number of degrees at which the individual images were taken and thus in the number of image captures (group A at 0.7° and group B at 1.0°, see Table 1). Hereby, the highest radiation dose in each measurement setting group was always measured in the three TLDs on the left thoracic side (red circles in Figure 2) and the lowest dose in the three TLDs on the right thoracic side (grey circles in Figure 2). Values of the ventral and dorsal side lay between the left and right sided TLD-measured radiation doses (black circles in Figure 2). The number of rotation steps was linear and had a proportional influence on the radiation dose. Thus, an increase of rotation steps from 0.7° to 1° led to a significantly lower radiation dose at the left thoracic side in the 1.0° measurement setting group compared to the 0.7° group for all analyzed filters (red columns in Figure 2, p<0.05).

**Figure 2:**
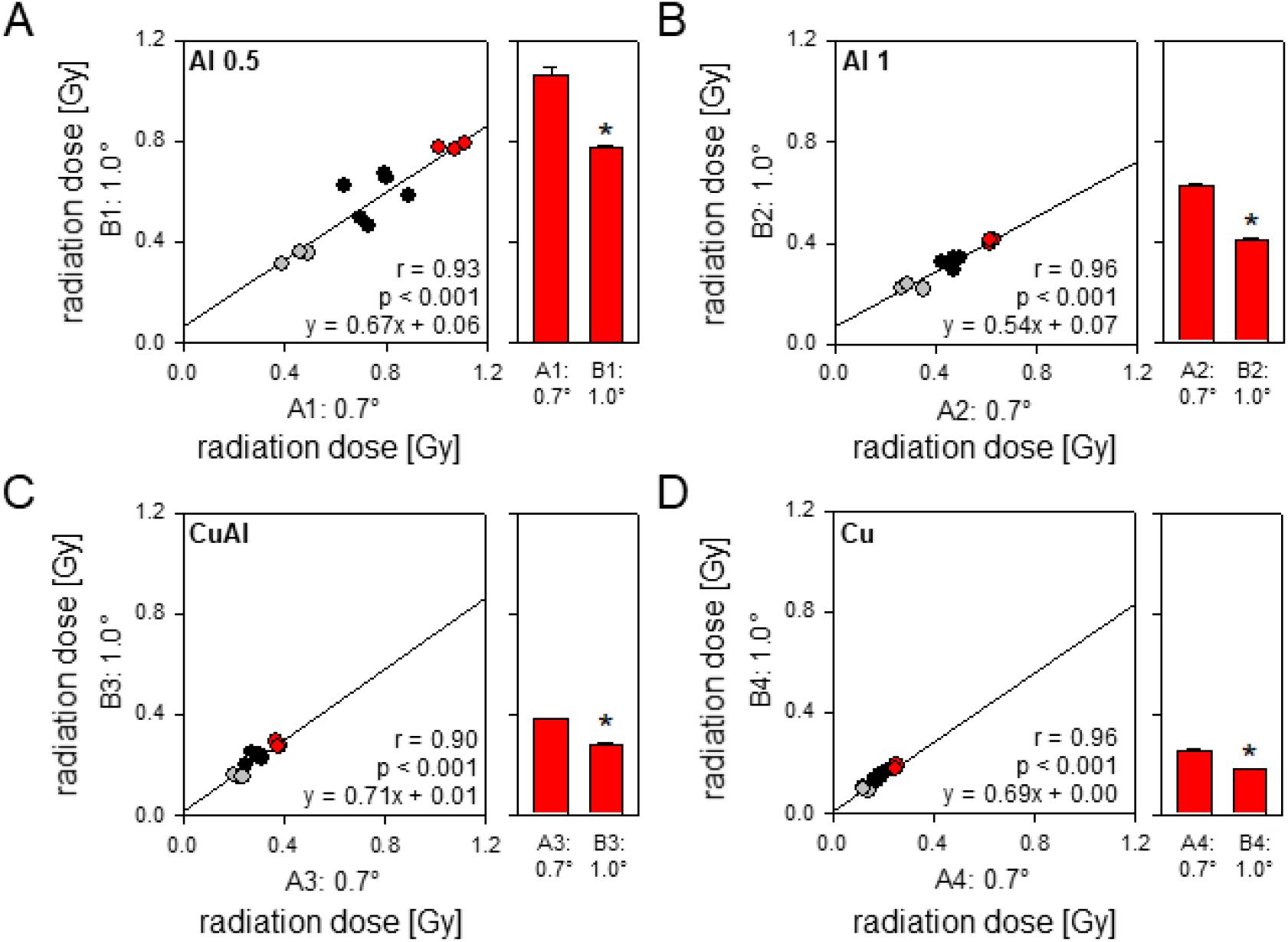
Impact of rotation steps on radiation burden. Image acquisition was performed using 0.7° (group A1-4) and 1.0° (group B1-4) rotation steps with a resolution of 35 µm. For all measurements, four different filters were used: Al 0.5 (**A**), Al 1.0 (**B**), CuAl (**C**), and Cu (**D**). A linear regression analysis was performed to evaluate the correspondence of each tested rotation step size using the same filter material. The radiation dose was measured using three TLDs each at the ventral (black), left (red), dorsal (black) and right thoracic side (grey) of the cadaver. The mean radiation dose of the left thoracic side is shown for each used filter. Means ± SEM. *p < 0.05.

### Impact of the resolution on the radiation burden

In a second set of experiments, micro-CT images were recorded at three different resolution levels: 35 µm for standard resolution (group B), 18 µm for medium resolution (group C) and 9 µm for high resolution (group D) in combination with each of the aforementioned filters. Hereby, 1.0° rotation steps were used for all applied resolution levels. During the measurements, the radiation dose at the left thoracic side (red circles in Figure 3) of the cadaver was markedly higher than the measured dose at the right thoracic side (grey circles in Figure 3). The radiation doses at the ventral and dorsal positions lay between those of the left and right thoracic side doses (black circles in Figure 3). Doubling the resolution from 18 µm (group C) to 35 µm (group B) reduced the radiation dose to 39-47 % of the original dose dependent on the selected filter (see slope of the correlation line) (Figure 3 A-D; p<0.001). For the resolution reduction by a factor of four from the highest tested resolution of 9 µm (group D) to the lowest tested resolution of 35 µm (group B) the radiation dose was reduced to 10-14 % of the original dose dependent on the selected filter (see slope of the correlation line; Figure 3 E-H; p<0.001).

**Figure 3:**
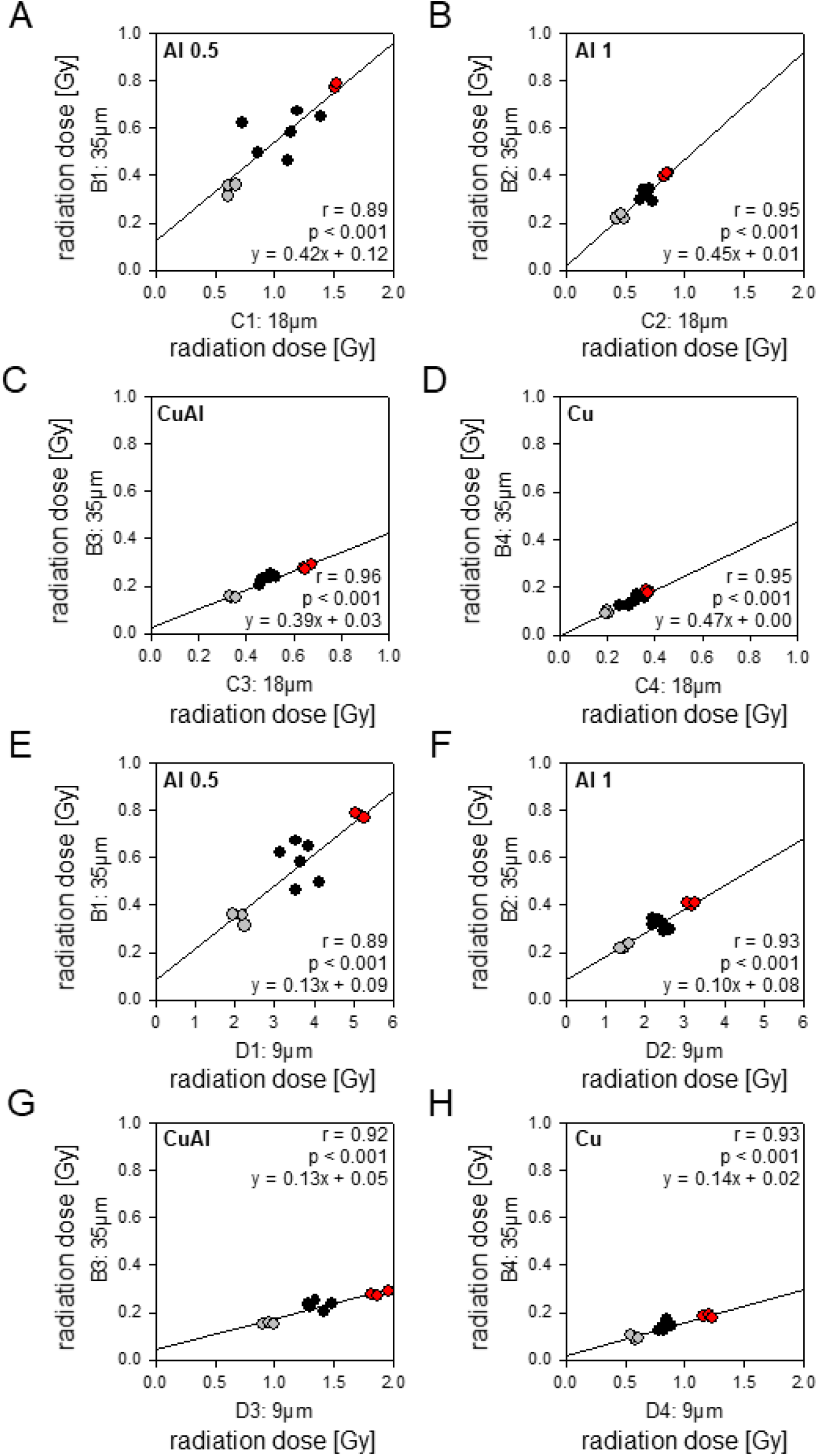
Impact of slice thickness on radiation burden. Image acquisition was performed using 35 µm (group B1-4; standard), 18 µm (group C1-4; medium) and 9 µm (group D1-4; high) for image resolution. Images of each group of resolution were performed using Al 0.5 (**A**+**E**), Al 1.0 (**B**+**F**), CuAl (**C**+**G**), and Cu (**D**+**H**) as available filters. Rotation step size remained in all three groups at 1.0°. The radiation dose was measured using three TLDs each at the ventral (black), left (red), dorsal (black) and right thoracic side (grey) of the cadaver. A linear regression analysis was performed to evaluate the correspondence of high and medium resolution compared to standard resolution using the same filter material. The slope of the correlation line indicates the reduction level of radiation dose dependent on the used filter. p<0.001

### Impact of the filter material on the radiation burden

Next, the dependency of radiation burden on the used filter materials and their thicknesses was examined. For this purpose, micro-CT image acquisition was performed with four different filters (Al 0.5, Al 1, CuAl, Cu) as described above. The Al 0.5 filter was compared to all other used filters. In general, both Al filters, i.e. Al 0.5 and Al 1, allowed a higher amount of radiation to pass than the CuAl and Cu filters (Figure 4A). Nevertheless, doubling of the Al filter thickness from 0.5 mm to 1 mm showed a significant reduction in radiation burden between 39 % at a resolution of 9 µm and 48 % at a resolution of 35 µm measured at the left thoracic TLDs (p<0.001). By using CuAl as filter material, the radiation burden was significantly reduced between 57 % at a resolution of 18 µm to 64 % at a resolution of 35 µm when compared to Al 0.5 (p<0.001). The highest radiation shielding was achieved by utilizing the Cu filter. Here, a reduction between 75 % at a resolution of 18 µm and 77 % at a resolution of 35 µm was detected (p<0.001).

**Figure 4:**
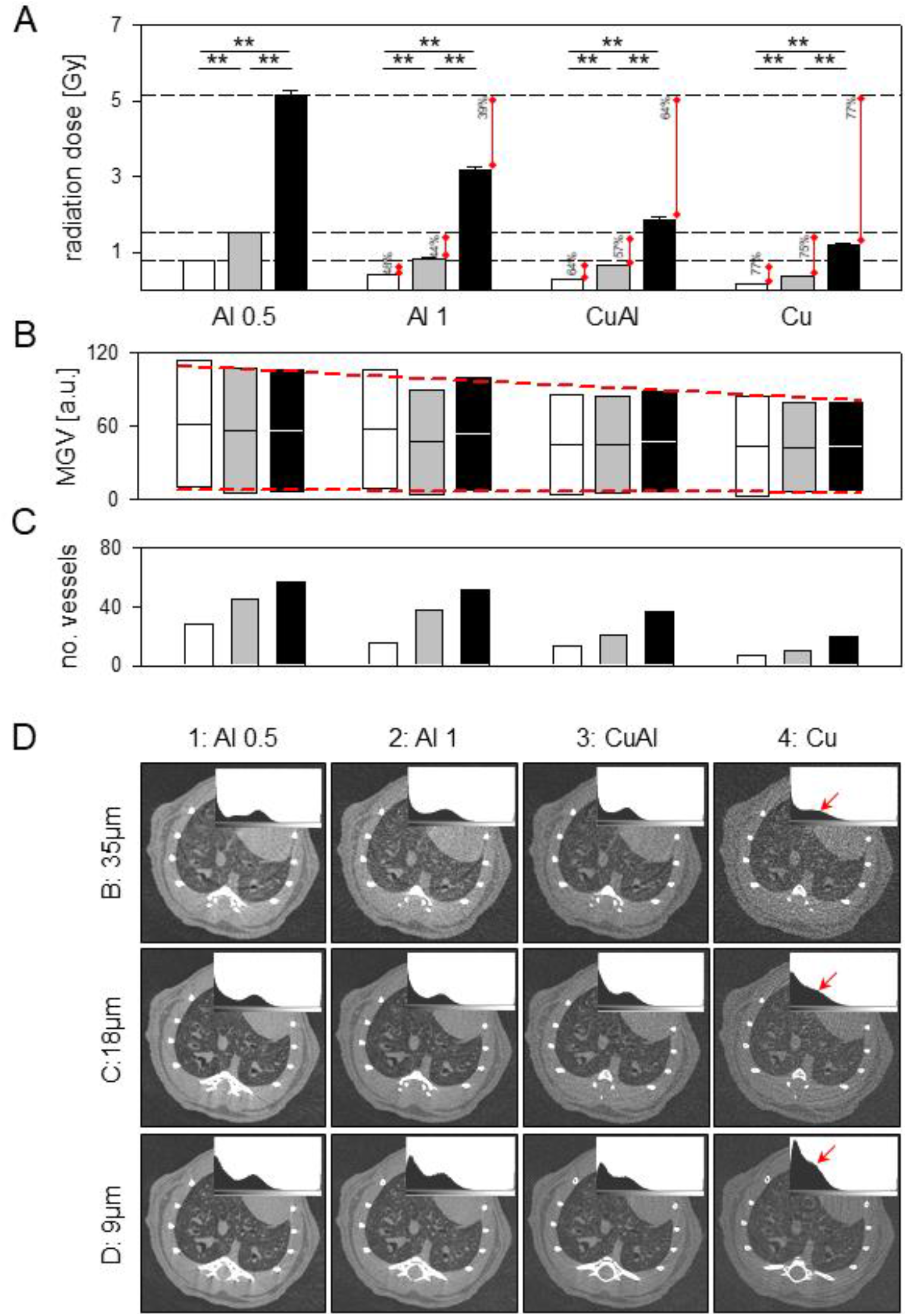
Impact of different filters on radiation burden. **A:** The radiation dose for the left thoracic side was analyzed for Al 0.5, Al 1, CuAl, and Cu in dependence of resolution (35 µm; 18 µm; 9 µm). Dashed lines indicate the level of left thoracic side radiation dose for 35 µm, 18 µm and 9 µm resolution imaged with Al 0.5. Means ± SEM. **p<0.001. **B:** The mean grey value (MGV) distribution in dependance of the used filter material and resolution is shown. Means ± SEM. **C:** Manually counted number of lung vessels in a micro-CT-slice at level of thoracic vertebra T4. As vessel count was performed only on one slice, no error bars are indicated. **D**: Representative images of transversal micro-CT scans at level of thoracic vertebra T4 using different resolutions (35 µm-B; 18 µm-C; 9 µm-D) and four different filter materials as indicated. The small boxes in each scanning condition show the histogram of grey values. Images acquired with Cu show flattening of the histogram scattering (red arrow). In **A** to **C**, white bars indicate 35 µm of resolution, grey bars indicate 18 µm of resolution, and black bars indicate 9 µm resolution level. The filter material is in the same order starting with Al 0.5 at the left over Al1, CuAl ending with Cu at the right position.

### Effect of the resolution and filter material on the image quality

Finally, the impact of the resolution and utilized filter material on the image quality of the micro-CT scans was examined. When analyzing the quality of the micro-CT images the largest mean grey value (MGV) distribution was measured in combination with Al filters, especially with Al 0.5 (Figure 4B). The MGV distribution measured exemplarily in all acquisition settings decreased from Al 0.5 over Al 1, CuAl until the smallest MGV distribution was achieved using the Cu filter (Figure 4B). Within the same filter material, no significant differences in MGV distribution was visible for different resolutions (Figure 4B).

To illustrate the influence of different image qualities as a consequence of resolution and filter material on the interpretation of scans, the number of lung vessels was manually counted on a slide at the level of thoracic vertebra T4 for every applied scan setting. The best image quality was achieved as a consequence of high photon density, resulting in a high MGV and a high number of identifiable lung vessels (Figure 4B-D). The highest numbers of lung vessels were detected at the highest resolution level of 9 µm in combination with Al 0.5. The number of detectable vessels decreased in dependence of the used filter material. Therefore, using Cu at a resolution of 35 µm showed the worst results in identifiable lung vessels combined with a low image quality, because of worse contrasting of different tissues (Figure 4C-D). The loss of contrast and MGV is indicated by the red arrow in Figure 4D.

## Discussion

In this study, we evaluated the impact of different micro-CT scanning modalities including filter material, rotation step size, and resolution on the obtained image quality and the overall radiation dose for mice. These factors are of a high impact for animal welfare in planning animal studies utilizing the micro-CT technology, particularly in case of long-term studies with repeated scans. We investigated four different radiation filters in combination with a scanning resolution of 35 µm, 18 µm, and 9 µm. Moreover, we tested the impact of different rotation steps on the overall radiation dose. In general, all micro-CT scans were performed counterclockwise over 180°. To measure the overall delivered radiation dose in each micro-CT protocol, we used a mouse cadaver, which was equipped with three TLDs each on the ventral, left, dorsal, and right thoracic wall ^21^. Best image quality especially for quantitative evaluation could be obtained by scanning at a high resolution level of 9 µm in combination with an Al 0.5 filter. However, this setting also generated the highest overall radiation dose, especially at the left side of the cadaver with an overall left-side dose of ∼5 Gy per scan. This total radiation burden could be minimized by using different filter materials as well as by increasing the rotation steps from 0.7° up to 1° which also minimizes the overall scan duration.Our results show that an increase of 30 % in rotation steps from 0.7° to 1.0°, accompanied by a simultaneous reduction of the scanning duration, is able to reduce the measured radiation dose significantly. This finding is in line with several studies that described a critical role of the scan duration in dependence of the obtained overall radiation burden ^8,12,14^.

In our measurements TLDs located on the right thoracic wall showed always the lowest radiation dose in all scan settings followed by TLDs located on the dorsal and ventral side. TLDs that were located on the left thoracic wall constantly displayed the highest measured radiation levels in every scan protocol independent of the selected filter and resolution. Here, it should be mentioned that the standard deviations were above the 5 % accuracy, as the TLDs could not always be exactly positioned in the same place for individual measurements. The TLDs were shrink-wrapped in small sachets to protect them from contaminations, which in turn could have a small impact on radiation dispersion. In addition, due to the small mouse body and the narrow radiation field there was an increased range of variation as compared to human measurements. As already mentioned above the radiation burden was divided to the same pattern of the local-specific TLDs in every examined micro-CT scanning protocol. This circumstance can be explained by the property of the scan protocol as well as by the characteristics of the scanner itself. The scan process runs over 180° starting from the dorsal position over the left ending at ventral position. Here, the left thoracic side is nearly constant in the primary radiation field and also closer positioned to the radiation source as any other thoracic side. Accordingly, TLDs that were located on the left thoracic side were exposed almost over the full scan duration, whereas the ventral and dorsal sides were exposed for a relevantly shorter time span. TLDs located on the right thoracic side were only exposed to transmitted and secondary radiation and, thus, exhibited a minimum of radiation doses.

In case of the used filters, it should be considered that different materials have different attenuation properties depending on their atomic number and mass density ^22,23^. Hence, Al (Z = 13) and Cu (Z = 29) show different attenuation patterns for X-ray photons also in dependence of their thickness and combination with each other ^24^. In 1988, Jennings reported that Cu filters are preferable, because Cu exhibits ∼10 % better shielding capacities when compared to Al ^25^. Nevertheless, Al is the most frequently utilized filter material. Using Al in different thicknesses, we could show that doubling the thickness from 0.5 mm to 1 mm achieved a reduction in radiation burden of 39 % up to 48 %. This reduction in radiation dose is a result of proportionally more low-energy radiation being attenuated by doubling the filter thickness. Cu is often used in pediatric radiology, because here it is most important to reduce radiation burden as much as possible. When using Cu as filter material in our study, radiation burden was even reduced up to 77 % compared to Al 0.5, which was the highest reduction we were able to generate by our settings. Due to its higher atomic number compared to Al, Cu reduces the entrance surface dose as well as the effective dose. However, in consequence, the obtained image quality is reduced due to a decrease in image contrast ^26,27^. In general, filtration of the X-ray beam minimizes the X-ray quantity in case of intensity reduction as described above for Cu. Here, the beam spectrum is varied by reducing low-energy X-rays and therefore beam quality. Due to the change of the radiation spectrum, the maximum beam energy is not minimized but maximized and thus the penetration depth of the beam is also increased ^15,28^. In this hardened spectrum, low-energy radiation is eliminated, which would be completely absorbed in the body and hence contribute to the total dose without yielding additional imaging information.

In some animal studies, e.g. focusing on tumor and/or metastatic growth over time or testing the effectiveness of new therapeutic approaches, it is necessary to perform serial micro-CT image acquisitions as a follow up. In these cases, the overall dose at the end of the experiment per animal must be estimated in advance to avoid cumulative radiation damage over time. Using a micro-CT protocol with an Al 0.5 filter, a rotation step of 1° and a resolution of 9 µm in our study, the obtained overall dose of 5.18 Gy was already higher than the LD50/30 of 5.0-7.6 Gy which is reported in the literature ^14,29^. Willekens et al. (2010) used a similar micro-CT setup in order to evaluate and optimize scanning protocols. In their study, doses between 0.295-0.507 Gy/scan were measured ^14^. The obtained doses were the highest in the subcutaneous tissue of the animals (6.81 mGy/mAs). Even in this dose range they postulated an influence on the experimental outcome, because they observed an inverse correlation between the survival time of the mice and the cumulative body dose ^14^. For whole body exposure a life shortening effect is quantified with ∼7.2 % per Gy, but depending on animal age, gender, tissue of interest, imaging time points and especially on the used mouse strain radiation tolerance varies and mice are able to recover after low radiation doses ^29,30^. Of interest, immune competent mice have a markedly higher ability to repair radiation damage in comparison to immunocompromised mice ^31,32^. Some mouse strains are able to repair irradiation damages directly within the initial minutes post radiation ^33^. Nevertheless, accumulated radiation-induced tissue damage can cause distress and pain and finally lead to animal death.

Although the studies described and discussed above differ significantly in some study design aspects and the used devices, one common point is the recommendation to keep the radiation dose as low as possible. This goal can only be achieved by accepting limitations in the obtained image quality. In our study the best image quality was achieved with an Al 0.5 filter in combination with the highest resolution level of 9 µm, leading by far to the highest radiation dose. Especially for long-term studies including repeated micro-CT scans, high image quality cannot be maintained, since the radiation exposure for the animals will be much too high. In these cases, we recommend to use alternative imaging modalities, such as optical imaging (OI), magnetic resonance imaging (MRI), positron emission tomography (PET), or single-photon emission computed tomography (SPECT) ^7^. For serial micro-CT acquisitions, we would recommend a scanner setting using Al 1 and a resolution of 18 µm, resulting in a dose of <1 Gy and one high-dose acquisition using Al 0.5 in combination with a resolution of 9 µm at the sacrifice day. Especially, if repeated micro-CT images are inevitable for a high validity of the experiment, our study provides valuable settings and a technical data background to minimize radiation exposure for the animals in consideration of the 3R principles and thus can help to optimize the planning of imaging modalities in longitudinal animal studies.

## Acknowledgements

The worthful contribution and excellent technical assistance of the animal keepers of the Institute for Clinical and Experimental Surgery is gratefully acknowledged.

## Research Data availability statement

Interested parties can contact the corresponding author to request access to the data.

